# A Surface-based deep learning approach for cortical shape analysis

**DOI:** 10.1101/2024.10.29.620757

**Authors:** Yanghee Im, Yuji Zhao, Boris A. Gutman, Sophia I. Thomopoulos, Elizabeth Haddad, Alyssa H. Zhu, Neda Jahanshad, Paul M. Thompson, Christopher R. K. Ching

**Affiliations:** Imaging Genetics Center, Mark and Mary Stevens Neuroimaging and Informatics Institute, Keck School of Medicine, University of Southern California, Marina del Rey, CA, United States; Department of Biomedical Engineering, Illinois Institute of Technology, Chicago, IL 60616, United State

**Keywords:** deep learning, cortical shape analysis, magnetic resonance imaging, UK Biobank, Alzheimer’s Disease, ADNI

## Abstract

Advances in deep learning hold promise for predicting clinical factors from human brain images. In this study, we applied a spherical harmonics-based convolutional neural network approach (SPHARM-Net) to MRI-derived brain shape metrics to predict age, sex, and Alzheimer’s disease (AD) diagnosis. MRI-derived brain features included vertex-wise cortical curvature, convexity, thickness, and surface area. SPHARM-Net performs convolutions using the spherical harmonic transforms, eliminating the need to explicitly define neighborhood size, and achieving rotational equivariance. Sex classification and age regression were carried out in a large sample of healthy adults (UK Biobank; N=32,979), and AD classification performance was tested in a large, publicly available sample (ADNI; N=1,213). SPHARM-Net showed strong performance for sex classification (accuracy=0.91; balanced accuracy= 0.91; AUC=0.97), and age regression (average absolute error=2.97 years; R-squared=0.77; Pearson’s coefficient=0.9). AD classification also performed well (accuracy=0.86; balanced accuracy=0.83; AUC=0.9). Our experiments demonstrate promising preliminary performance using the SPHARM-Net for two widely studied benchmarking tasks and for AD classification. Future work will include comparisons of shape-based methods and extending these analysis to more challenging tasks such as mood disorder classification.

## 1. INTRODUCTION

Surface-based methods have been used to understand focal variations in brain morphometry associated with age, sex and the presence or progression of neurological conditions such as Alzheimer’s disease (AD). Surface-based shape features can be generated from structural MRI using a variety of methods, and often use 3D triangulated meshes to calculate vertex-wise measures across thousands of points across the brain. These include cortical thickness, surface area, curvature, and convexity.^1,2^ Prior studies show brain surface area differences in men and women.^3^ Advancing age is associated with thinner cortex and surface area, with more severe patterns observed in AD.^4^ However, the extent to which structural brain shape measures could be used to predict clinical decisions like distinguishing healthy agers from those who will go on to develop AD, remains unclear.^5^

Convolutional neural networks (CNN) have been widely applied to structural brain MRI features. Applying CNNs to cortical surface data remains challenging as brain surface models are generally represented as irregular grids. Two general approaches have been evaluated for surface-based deep learning using vertex-wise brain shape measures: (1) graph convolutional neural networks (GCN), and (2) spherical convolutional neural networks (spherical CNN). GCNs have been used for surface-based analysis for age prediction, sex classification, and AD classification.^6–8^ To apply GCNs to brain surface data, surfaces must be registered into a common space to establish vertex-wise correspondence across individuals. This can compensate for distortions in the sulcal pattern resulting from non-rigid surface registration, which is designed to improve the alignment of cortical folds. 1-ring or 2-ring neighborhoods are also often used to enhance computational efficiency. A recent approach from Liu et al. proposed a vertex-convolutional neural network (vertex-CNN), which consists of convolution and pooling operations, based on a GCN with a 1-ring neighborhood on the vertex domain.^9^ This method performed well when classifying individuals with mild cognitive impairment and AD using the ADNI dataset, and also showed superior computational efficiency compared to GCN. Despite the increased efficiency, GCN performance is highly dependent on spatial sampling factors such as graph regularity and edge length. Spherical CNNs address these challenges by operating in the spherical domain. Common approaches to convolve on the sphere include spatial sampling along a spherical grid and spectral transformation. One effective way is to re-tessellate the unit sphere using an approximately uniform grid, such as icosahedral subdivision,^10^ for an approximation of the solution as a sum over the spherical grid. Despite these strategies, the finite nature of spherical grids poses challenges for rotation-invariant convolution, which highlights the need for more robust spherical convolution methods that can handle these constraints effectively.

In this study, we extend the SPHARM-Net approach to predict demographic factors including age, sex, and AD diagnosis. The first two of these tasks are common benchmarking tasks where the ground truth is known, and the third diagnostic task could have value in clinical settings. We first assessed whether the model could learn relevant features by performing age regression and sex classification in a large cohort of healthy adults (N=32,979). Then we evaluated its performance for AD diagnosis classification using accuracy, balanced accuracy, and a receiver operating characteristic area under the curve (AUC).

## 2. METHODS

### Datasets

In this study, 3D T1-weighted (T1w) brain MRIs from healthy control (HC) participants were used to perform sex classification and age regression in 32,979 participants from the UK Biobank study (age ranged 44.6-82.8; mean age: 64.0 yrs. ± 7.7SD; F=17,314/M=15,665). HC participants were derived using the UKB Data Parser^11^ and defined as having no ICD-10 history of illness with known effects on brain structure. For AD classification, brain MRI were used from 379 individuals with AD and 834 HC (N=1,213; age ranged 55.1-91.0; mean age: 73.4 years ± 6.9 SD; F=639 /M=576) from the Alzheimer’s Disease Neuroimaging Initiative (ADNI; adni.loni.usc.edu).

### Cortical Features

FreeSurfer (v7.1.1.) was used to reconstruct vertex-wise cortical shape features across 163,842 vertices. We derived the geometric features at each vertex: mean curvature of inflated and smoothed gray/white surfaces, average convexity, cortical thickness, and surface area. Each participant’s sphere was re-tessellated using an icosahedral subdivision of 40,962 vertices.

### Spherical Harmonics-Based Convolutional Neural Network (SPHARM-Net)

SPHARM-Net^12^ performs convolutions using the spherical harmonic transform, eliminating the need to explicitly define neighborhood size and achieving rotational equivariance. This rotationally equivariant network handles anatomical variability without extensive data augmentation or rigid alignment. A key innovation here is the constrained spherical convolutional filter, which supports an infinite set of spectral components, avoiding harmonic truncation and preserving geometric details.

### Network Architecture

We adapted SPHARM-Net to implement classification/regression by excluding the decoder portion of the network. Our model consisted of a full bandwidth spherical harmonics-based spectral convolutional layer, three spherical harmonics-based spectral convolutional layers, a global average pooling layer, and a fully connected (FC) layer that outputs the predicted clinical factor. Each spectral convolutional layer has a spherical harmonic transform, its inverse spherical harmonic transform, a convolutional layer between them, a batch normalization layer, and a non-linear rectified linear unit activation function. Spectral pooling layers were used after each spectral convolutional block to limit the bandwidth of the convolution without adjusting the spatial resolution. Each block reduces the harmonic bandwidth *L* by half while decreasing the number of channels *C* by a factor of two.

### Experiments

To evaluate the capacity of SPHARM-Net to learn meaningful features from brain MRI shape features, we first conducted sex classification and age regression (two widely studied benchmarking tasks). Then, we trained a model for AD classification in a separate dataset. Data were split into independent training, validation, and testing sets, approximately 70:20:10, respectively. For each task, we performed a grid search to select hyperparameter values, including the learning rate {1e-4 to 1e-6}, optimizer {ADAM, ADAM with weight decay}, encoder layers {2, 3, 4}, and bandwidth {80, 120} using 5-fold cross-validation. SPHARM-Net was trained for 30 epochs for each scan using the Adam (with weight decay) optimizer with a learning rate of 1e-4 or 1e-6 and decay by a factor of 0.1 if no improvement was made in two consecutive epochs. Through the extensive hyperparameter search, cortical thickness and surface area were chosen as inputs to the model for age regression, while curvature and convexity were used for sex classification and AD classification. Final model selection was based on the lowest validation loss. For regression, the model was evaluated using mean absolute error (MAE), R-squared (R^2), and the correlation coefficient (R). For classification tasks, accuracy (ACC), balanced accuracy (BAC), and AUC were used for the evaluation of the model.

## 3. RESULTS

**Table 2** shows the results obtained for age regression, sex classification, and AD classification. The model showed an accuracy of 0.91, a balanced accuracy of 0.91 and AUC of 0.97 for sex classification, an average absolute error of 2.97 years, a R-squared value of 0.77, and a Pearson’s coefficient of correlation of 0.9 with for age regression, and an accuracy of 0.860, balanced accuracy of 0.83 and AUC of 0.9 for AD classification. **Figure 1** shows the scatterplot of the predicted age chronological age for the test set of UKB. Receiver operating characteristic curves of the model for sex classification and AD classification are illustrated in **Figure 2**.

**Table 1.** Data distributions for independent training, validation, and test sets.

| UKB (N=32,939) | N | Age range (mean ± SD) | Sex (F/M) |
| --- | --- | --- | --- |
| Training | 19,788 | 44.6-82.8 (64.0±7.7) | 10354/9434 |
| Validation | 6,595 | 46.1-82.3 (63.9±7.7) | 3489/3106 |
| Test | 6,596 | 45.2-82.4 (64.1±7.7) | 3471/3125 |
| ADNI (N=1,213) | N | Age range (mean ± SD) | Sex (F/M) |
| Training | 776 | 55.3-91.0 (73.6 ± 6.9) | 411/365 |
| Validation | 194 | 56.0-90.3 (73.4 ± 7.0) | 92/102 |
| Test | 243 | 55.1-89.9 (72.9 ± 6.8) | 135/108 |

**Table 2.** Results of age regression, sex classification, and AD classification. Abbreviations - MAE, mean absolute error; R^2, R-squared value; R, Pearson correlation; ACC, accuracy; BAC balanced accuracy; AUC, a receiver operating characteristic area under the curve; CI, confidence interval.

| Task | Prediction | Features | MAE | $R^2$ | R |
| --- | --- | --- | --- | --- | --- |
| Regression | Age | Thickness + Area | 2.97 | 0.77 | 0.9 |
| Task | Prediction | Features | ACC | BAC | AUC (95% CI) |
| classification | Sex | Curvature + Convexity | 0.91 | 0.91 | 0.97 (0.97-0.98) |
| classification | AD | Curvature + Convexity | 0.86 | 0.83 | 0.9 (0.85-0.94) |

**Figure 1.**
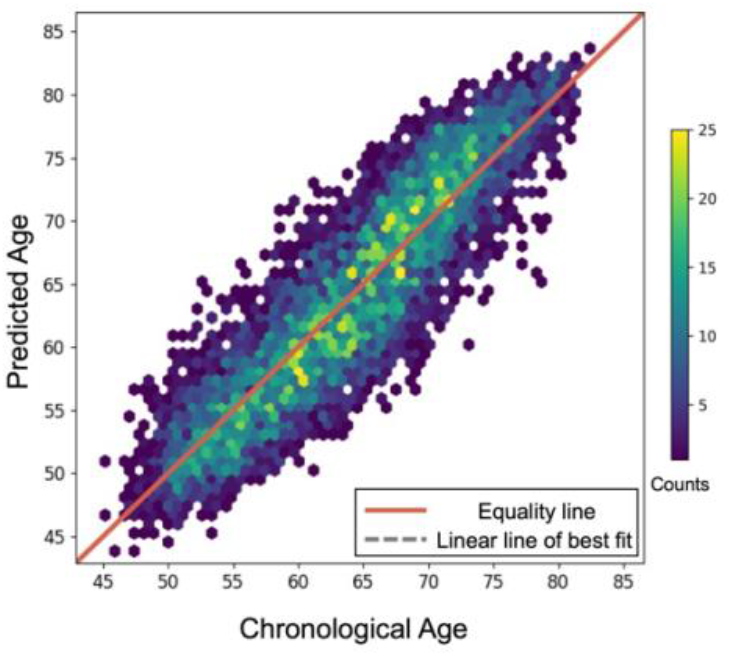
Scatterplot of the chronological age and predicted age from the test set (N=6,596). Each dot represents one individual. The color of the dots indicates the number of individuals in that position.

**Figure 2.**
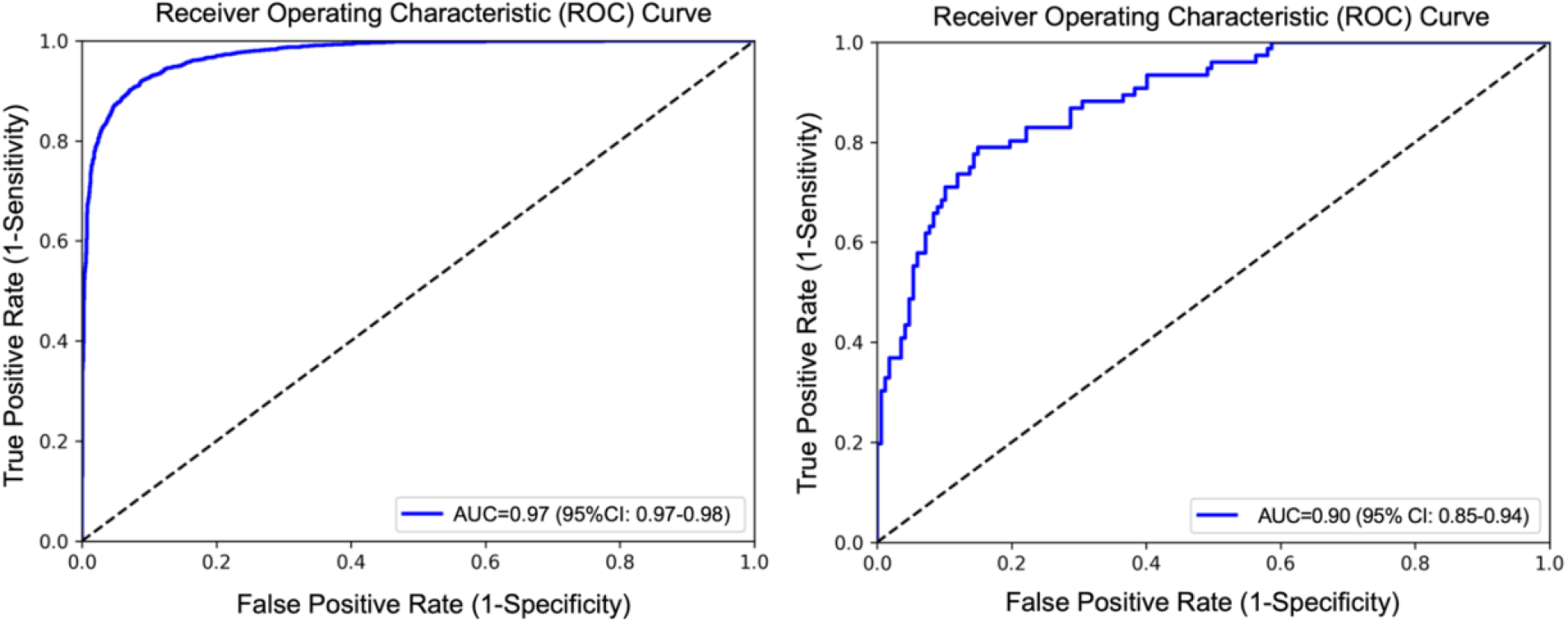
Receiver operating characteristic curve of the model for sex classification (left) and AD classification (right) using curvature and sulcal depth.

## 4. CONCLUSION

Here we adapt a surface-based deep learning method using large-scale MRI data for various prediction and regression tasks. This analysis pipeline offers a roadmap for future large-scale deep learning studies of brain imaging beyond age regression, sex classification, and Alzheimer’s disease classification. In the future, we will adapt this approach to include subcortical surface-based shape features and more challenging tasks such as identifying individuals with mild cognitive impairment or mood disorders. These experiments could provide insights into the generalizability of these models relative to other surface-based techniques.

## ACKNOWLEDGEMENTS

This work was supported by NIH grants R01 MH121806 and R01 MH129742.

